# Temporal changes in mechanical pin prick sensitivity following high frequency induced sensitisation of central nociceptive pathways: a test re-test reliability study

**DOI:** 10.64898/2026.01.09.698631

**Authors:** Samuel Mugglestone, Giorgio Ganis, Sam W. Hughes

## Abstract

High-frequency stimulation (HFS) is a human surrogate model of secondary hyperalgesia and a key experimental tool for understanding the mechanisms and modulation of central nociceptive pathways. An emerging area of research focuses on the role of top-down endogenous analgesic systems during secondary hyperalgesia development. However, the test-retest reliability of the early temporal changes in sensitivity are poorly understood. In the present study, we investigated the between-session reliability of the early temporal dynamics and late-phase expression of HFS-induced changes in mechanical pinprick sensitivity in a heterotopic area on the volar forearm in 28 healthy participants across five time points relative to HFS conditioning: −15, 5, 20, 35, and 50 minutes. Homotopic changes in single-pulse electrically evoked responses were also assessed although no primary hyperalgesia was evident. Baseline conditioned pain modulation (CPM), temporal summation of pain (TSP), and state-trait anxiety (STAI) were also assessed to investigate potential influences on heterotopic and homotopic responses. The present findings demonstrate the consistent induction of mechanical pinprick secondary hyperalgesia by the end of the HFS window (50 minutes) across repeated test session. However, a distinct reduction in the development of sensitivity was present during session 2. Furthermore, pain during HFS conditioning, anxiety, CPM, and TSP demonstrated no influence on secondary hyperalgesia development and were inadequate to explain between-session variance. These results suggest that careful planning around experimental designs, and the counterbalancing of experimental conditions should be considered when investigating modulating factors over the development of secondary hyperalgesia. Further research into factors influencing habituation across sessions is needed.

**Perspective:** High-Frequency Stimulation evokes mechanical secondary hyperalgesia across repeated sessions; however, sensitivity development is diminished, and unexplained by pain intensity during HFS conditioning, anxiety, or cuff algometry CPM and TSP.

## Introduction

High-frequency stimulation (HFS) is a human surrogate model of secondary hyperalgesia (SHA) and is therefore a key tool for understanding mechanisms and modulation within central nociceptive pathways ^1–3^. HFS can induce homosynaptic and heterosynaptic plasticity in the spinal cord dorsal horn, manifesting as primary hyperalgesia (PHA) and secondary hyperalgesia (SHA) respectively ^2, 4^.

Punctuate mechanical pinprick sensitivity (MPS) is often used to characterise SHA through the activation of both high- and low-threshold cutaneous mechanoreceptors in the area adjacent to HFS conditioning ^1, 5, 6^. Within the HFS model, the presence of SHA follows a broad time course in which increased MPS is evident at 30 minutes and maintained over hours ^3, 4, 7^. Although some reports have also demonstrated that prominent SHA is present as early as 15 minutes ^1, 3^. It is therefore important to further investigate both the late phase expression of SHA alongside early temporal dynamics and whether these responses are consistent across repeated test sessions. Whilst test-retest reliability has been previously demonstrated at later time points, with scores ranging from moderate to good ^8, 9^, the reliability of early dynamic changes that occur during the progression of SHA are poorly understood.

Additionally, since the initial proposal of the HFS model, the development of homotopic facilitation remains underexplored and attempted replications applying homotopic single-pulse electrical stimuli have resulted in contradictory findings, with some reporting an increase and some reporting a decrease in pain ratings ^2, 4, 10, 11^. Therefore, it is crucial to investigate the consistency and reliability of EPP changes post-HFS within the same individuals across repeated test sessions.

Such electrical pain perception (EPP) is considered to be the result of selectively activating superficial A-delta and C fibre nociceptors, although this specificity may be dependent upon the intensity of stimulation ^1, 12, 13^.

Furthermore, recent investigations have demonstrated that HFS-induced changes in somatosensory function, can be manipulated by psychosocial factors ^14–17^. Specifically, experimental manipulations of anxiety-related states, such as pain expectations or the fear of pain, have been shown to influence the magnitude of SHA ^17^. Additionally, trait anxiety substantially alters pain perception within both healthy controls and clinical cohorts ^18^. Therefore, to support the use of HFS as a reliable experimental model it is important to clarify whether individual differences in anxiety change HFS-induced SHA.

In addition, descending pain control, as assessed through conditioned pain modulation, does not explain inter-individual variability in the development of HFS-induced SHA ^19, 20^. Therefore, sufficiently intense and prolonged activation of peripheral nociceptors may bypass inhibitory control ^20, 21^. However, the test-retest reliability of this finding and the role of endogenous inhibitory mechanisms in explaining between- and within-individual variability in HFS-induced SHA have yet to be investigated. Therefore, it is important to assess whether individual differences in CPM predict the development of SHA.

Temporal summation of pain (TSP) is often assessed in parallel to CPM, reflecting the facilitation of central ascending nociceptive pathways in the spinal cord. As opposed to HFS, TSP provides a measure of spinal excitability and does not induce longer-term synaptic modifications. However, the two experimental models may share overlapping mechanisms of facilitated nociceptive activity in the dorsal horn ^22^. Despite speculation of a predictive relationship between TSP and the magnitude of SHA ^20^, there is a lack of experimental evidence to demonstrate this association.

In the present study, the test-retest reliability of heterotopic changes in MPS and homotopic changes in single-pulse EPP across multiple time points following HFS conditioning was investigated. It was hypothesised that within both sessions, pronounced homotopic and heterotopic sensitivity would occur, as indexed by a gradual increase in pain ratings ^2, 4^. It was also anticipated that these changes would show good reliability ^9^. Additionally, individual differences in state and trait anxiety, CPM, and TSP, were hypothesised to contribute significantly to the magnitude of secondary hyperalgesia and the development time course, demonstrating either predictive or confounding relationships.

## Methods

### Pre-registration

This study is part of a wider research program investigating the reliability of behavioural and neurophysiological methods for assessing changes to central nociceptive function induced by HFS conditioning. The overall study protocol was preregistered on the Open Science Framework (OSF) and is available at: https://doi.org/10.17605/OSF.IO/S5A9Y

### Participants

Recruitment was conducted via the University of Plymouth’s Paid Participation Pool alongside word of mouth. Participants were instructed to abstain from alcohol and drugs prior to each visit, and to specifically avoid taking painkillers and to limit caffeine intake. At the start of each visit compliance was confirmed. All participants provided written informed consent and were informed of their continual right to withdraw. The study was approved by the University of Plymouth Ethics Committee (ID:3777) and in compliance with the Declaration of Helsinki (World Medical Association, 2013).

### Experimental design

Two functionally identical sessions were conducted at least a week apart with the aim of assessing the between-session reliability of the temporal changes in heterotopic and homotopic responses following HFS-conditioning. At the beginning of each session state-trait anxiety inventory (STAI), TSP, and CPM data were collected sequentially. Before beginning the main experimental procedures, participants were familiarised with the sensory testing stimuli and electrical detection threshold (EDT) was determined over the volar surface of the forearm. Baseline MPS and EPP responses were initially recorded on the ipsilateral and contralateral forearms, followed by HFS conditioning applied only to the ipsilateral forearm, not the control arm. MPS and EPP responses were then reassessed at 4 time points post-HFS with 15-minute intervals between the beginning of each block (Figure 1a-c).

**Figure 1.**
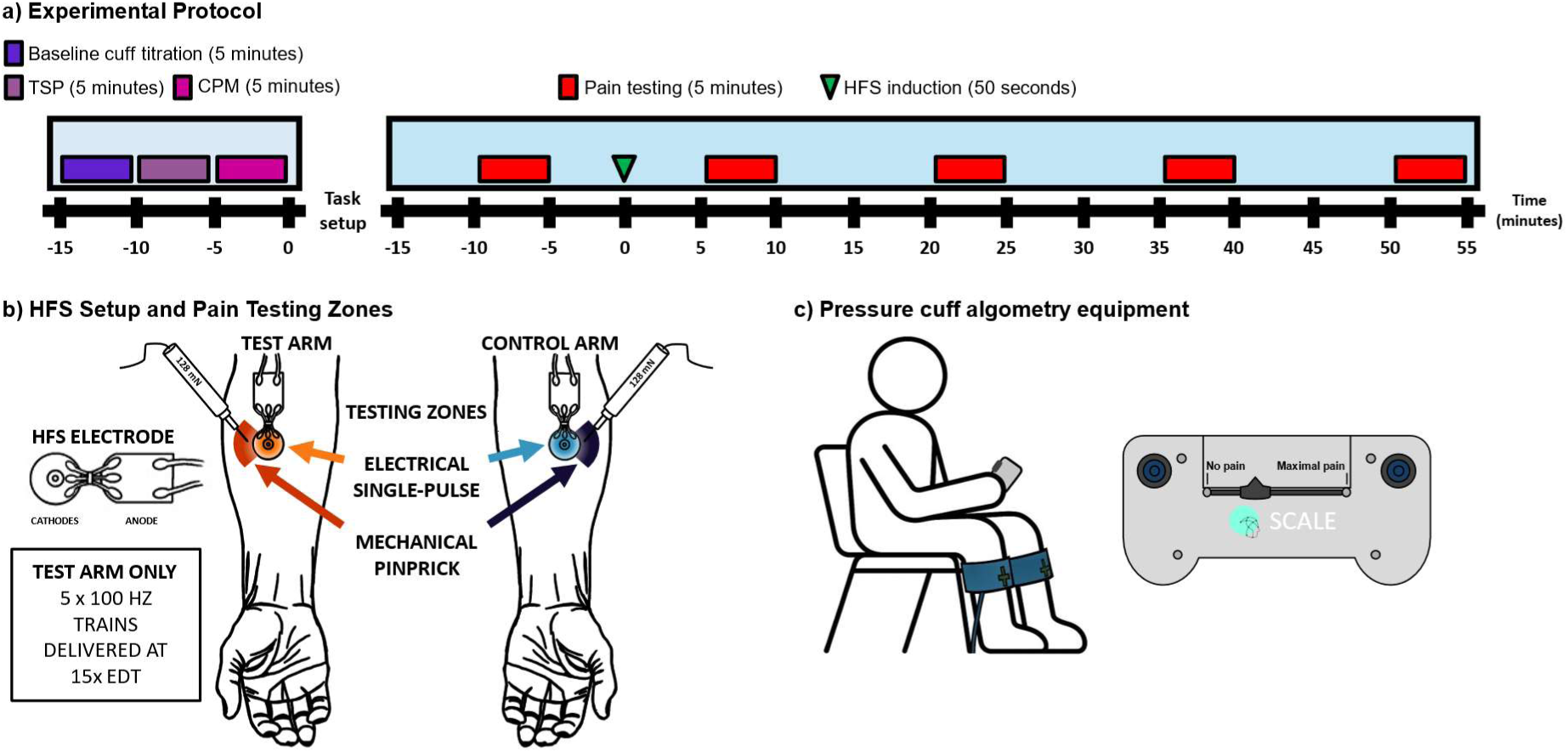
Experimental protocol. a) Identical experimental procedures were followed for both sessions. Baseline cuff titration = Ischemic cuff pressure thresholding; TSP = Temporal Summation of Pain; CPM = Conditioned Pain Modulation; Pain testing = A series of single-pulse electrical stimuli and mechanical pinprick stimuli, twenty of each type were applied to both the test and control arms in a pseudo-randomised order; HFS = high-frequency stimulation. b) Identical electrodes were placed on the volar forearms of each subject and remained throughout the session to ensure consistent delivery of single-pulse electrical stimuli to the homotopic zone. Individual EDT was established at the beginning of session 1 and carried over to session 2. All electrical stimuli were delivered at 15 times EDT. Mechanical pinpricks were administered to the heterotopic zone outside of the electrode site. All pinprick stimuli were 128 mN. HFS was administered exclusively to the test arm (right arm) in five trains of 100 Hz (15 x session 1 EDT). Subjects were familiarised with both stimuli modalities at the beginning of each session and required to report pain intensity ratings verbally via a numerical rating scale (0 – 100). EDT = Electrical detection threshold. c) Identical algometry cuffs were placed around the gastrocnemius muscles of each subject whilst participants were sitting. A visual analogue scale with slider and response buttons was used to collect pain measurements and provide control over the pressure shutoff. Baseline measurements were taken for each leg separately. Pain detection threshold was determined as the pressure corresponding to the first movement of the scale. Pain tolerance threshold as determined as the pressure corresponding to either response button being activation, or upon reaching the safety threshold.

Within each block, 20 stimuli were delivered per condition (arm x modality) with an interstimulus interval of ∼2 seconds resulting in 40 stimuli per arm, and a total of 80 stimuli within each time point as both mechanical pinprick and single-pulse electrical stimulation were assessed, albeit in separate zones. This was considered the minimum number of trials to effectively characterise the neurophysiological components of the wider research program. This remains within standard DFNS QST protocols for the assessment of MPS which involves collecting 35 pain ratings at a single time point ^23^. Comparable studies have ranged from 3 ^24^ to 160 ^25^ stimuli within a single block, with typical assessments between 20 and 30 stimuli per condition ^1, 26–32^.

### High-Frequency Stimulation

Cutaneous electrical stimuli were applied to the right volar forearm (5cm distal to the cubital fossa) using an EPS-P10 electrode (MRC Systems GmbH, Heidelberg, Germany) via a constant current stimulator (2ms pulse width; DS7; Digitimer Ltd; Welwyn Garden City, UK) (Figure 1b). Prior to attaching the HFS electrode, the skin was cleaned with a non-alcoholic aqueous solution and allowed to dry. The EPS-P10 electrode contains a circular array of 10 cathodal pins (diameter: 0.25mm) and a larger surface area anode (area: 410 mm2). Such concentric surface electrodes have been shown to selectively activate A-delta and C fibre free nerve endings located in the superficial epidermal layers without the excitation of A-beta fibre mechanoreceptors located in the dermis ^12, 33^. Sufficiently high intensity electrical stimuli can bypass peripheral transduction mechanisms directly activating voltage-gated ion channels ^1^. Notably, this specificity may falter depending upon the intensity of stimulation, resulting in the activation of deeper A-beta fibres ^13^. However, characterising the incidence of this phenomenon is beyond the scope of the current study.

Electrical detection threshold was determined using 2ms pulses starting at 0.05 mA in 0.05 mA increments until participants reported a clear sensation, then decreased by 0.01 mA until the sensation was lost. The intensity was increased in 0.01 mA increments until the sensation returned, repeating this process three times. The average of the six recorded intensities represented each participant’s EDT. Trains of HFS (100 Hz; NL304 Period Generator; Digitimer Ltd) were applied 5 times to the test (right arm) for 1 second each (NL405 Width Delay; Digitimer Ltd) at 15x EDT every 10 seconds (i.e. over a 50 second period). The participants were asked to the rate the intensity of the HFS after each train using a numerical rating scale (NRS; 0 = no pain, 100 = worst pain imaginable). Notably, whilst individual HFS stimulation intensity is typically re-thresholded per session, the present study carried over identical intensities for each participant to allow for a clearer evaluation of any habituation across sessions. Furthermore, given the variability in the development of SHA across sessions, habituation to HFS stimuli was assessed through changes in average pain intensity during HFS conditioning via intraclass correlation coefficients, linear-mixed effects modelling, and visual inspection of individual variability.

### Heterotopic mechanical stimulation

Mechanical pain sensitivity (MPS) was assessed across both forearms using a 128 mN pin prick stimulator within a typical heterotopic area laterally adjacent to the conditioning electrode yet within 2 cm as depicted in Figure 1b ^7, 34^. However, the area of SHA was not systematically mapped. Although mechanical pinprick stimuli activate both high- and low-threshold mechanoreceptors (including nociceptive A-delta and C fibres), low-intensity pinpricks can be perceived as non-painful ^1^. Therefore, to avoid floor or ceiling effects, the medium range intensity of 128mN was used throughout the study and baseline pain ratings are reported.

Following every five successive stimuli, participants rated their average perceived pain on a numerical rating scale (NRS; 0 - 100). These were further averaged to provide a single value for each time point (Figure 2a). Therefore, within each time point, participants received a total of 20 pinprick stimuli to each of the ipsilateral and contralateral arms.

**Figure 2.**
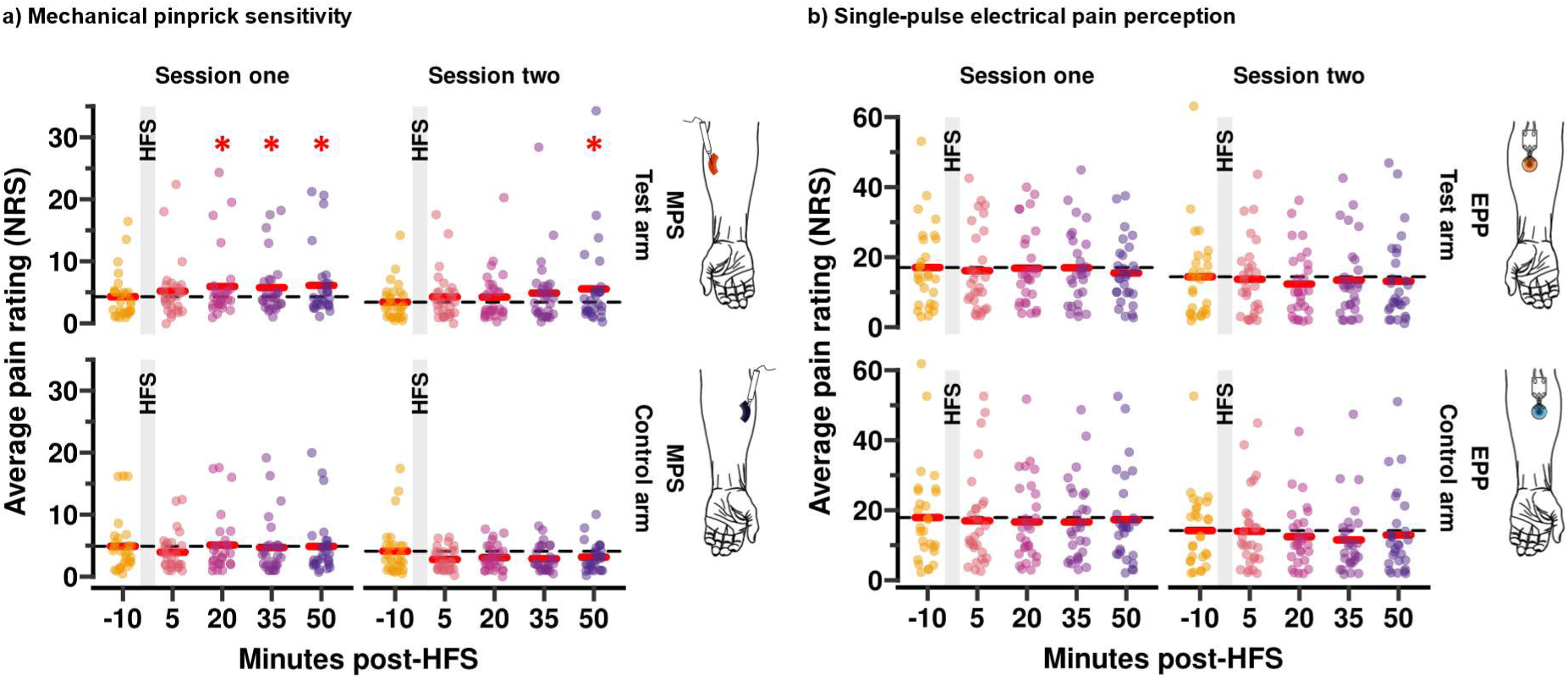
a) MPS and b) EPP results for sessions one (left) and two (right) for the test (upper) and control (lower) arms. Raw data points are displayed as an average for each individual at each time point, session, and arm. Red bars indicate group means. Statistics were derived from linear mixed-effects modelling and are comparative to the baseline of each arm respectively. * Indicates significance at alpha = 0.05; HFS = High-frequency stimulation; MPS = Mechanical pinprick sensitivity; EPP = Electrical pain perception; NRS = Numerical rating scores.

### Homotopic electrical stimulation

Electrical pain perception (EPP) was assessed across both forearms using the same epi-cutaneous pin electrode as used to deliver HFS, with the same intensity of 15x EDT as outlined above. However, each stimulus consisted of a single square wave pulse via a constant current stimulator (2ms pulse width; DS7; Digitimer Ltd; Welwyn Garden City, UK) which were delivered directly to the area of HFS conditioning. Both test- and control-arm electrode were positioned prior to baseline assessments and remained in place throughout the main experimental procedures 5cm distal to the cubital fossa as depicted in Figure 1b. Such stimulation can selectively activate A-delta and C fibres through bypassing peripheral transduction mechanisms and directly activating voltage-gated ion channels within superficial afferent nociceptors ^1^.

Following every five successive stimuli, participants rated their average perceived pain on a numerical rating scale (NRS; 0 - 100). These were further averaged to provide a single value for each time point (Figure 2b). Therefore, within each time point, participants received a total of 20 single-pulse electrical stimuli to each of the ipsilateral and contralateral arms.

### Cuff-algometry protocol

CPM and TSP protocols were conducted with computer-controlled cuff-algometry system (LabBench software 4.7.3 and CPAR+ device, Aalborg, Denmark). Independent computerised torniquet cuffs were placed around each calf (10 x 61 cm) ^35, 36^ (Figure 1c). Test stimulus and conditioning stimulus were always applied to the right and left calves, respectively. Participants were handed a computerised visual analogue scale (VAS) anchored from ‘no pain’ to ‘maximal pain’ and a slider set to ‘no pain’ as a default.

First, pain detection and tolerance thresholds were measured. The test stimulus cuff began to inflate at a rate of 1 kPa/sec to a maximum of 100 kPa. Participants were instructed to move the slider on the computerised VAS once they felt their first pain sensation and continuously rate their pain until the maximum they could tolerate. Once they reached their maximum tolerable pain, participants could push a ‘stop’ button placed next to the computerised VAS, which deflated the cuff immediately. The pressure at which participants first moved the VAS slider was considered their pain detection threshold (PDT), and the pressure registered immediately before participants pressed the ‘stop’ button was considered their pain tolerability threshold (PTT). If participants did not press the ‘stop’ button before reaching the maximum pressure, then this was considered their PTT (i.e., 100 kPa). Following this, an identical examination was performed on the conditioning stimulus cuff.

Following a 2-minute resting period, the TSP protocol was carried out on the test stimulus calf; ten 1-second stimuli with pressure set to participants individual PTT recorded during the earlier assessment on the same calf. The cuff inflated and deflated rapidly for each stimulation. Stimuli had a 1-second duration with 1-second interval between them. Participants were instructed to rate their perceived pain intensity on each stimulus using the computerised VAS without returning it to ‘no pain’ between stimuli to ensure stable measurements and avoid lurching movements.

Following a 5-minute resting period, the final part of the CPM protocol was carried out. The conditioning stimulus cuff (left calf) inflated rapidly to a pressure equal to 70% participants’ PTT on the same calf, and it remained inflated until the end of the protocol to serve as concurrent conditioning tonic stimulus. Once it reached the target pressure, the test stimulus cuff started to inflate at a rate of 1 kPa/sec (i.e. using a parallel CPM design). Participants were instructed to rate the test stimulus (right calf) in the exact same way as they did during thresholding.

#### CPM analysis

Within each session, two strategies to calculate CPM effects were followed. On one hand, CPM was assessed as a change in pain *thresholds* during tonic stimulation. CPM measures, calculated separately for PDT and PTT, resulted from the absolute difference in pain thresholds of the test stimulus during baseline and conditioning tests (i.e., CPM = pain threshold _baseline_ – pain threshold _conditioning_). Here, positive and negative CPM values reflected descending facilitation and inhibition, respectively. In addition, CPM was also assessed as a change in overall pain *ratings* throughout the trials. CPM measures resulted from the absolute difference in the area under the curve (AUC) for continuous VAS ratings of the test stimulus within the overlapping range across baseline and conditioning tests for each run (i.e., CPM = AUC _baseline_ – AUC _conditioning_).

Given the variability in CPM demonstrated through intra-class correlation coefficients (Supplementary Table 1), CPM responder rates were categorised based upon the direction and degree of change relative to standard error of measurement (SEM). SEM is calculated as the standard deviation of the baseline measure times the square root of 1 minus the intraclass correlation coefficient (3, 1): Values < 1 SEM are weak responders; values between 1 SEM and 2 SEM are responders; values > 2 SEM are strong responders ^37^. As is typical, not all participants exhibited a strong inhibitory CPM effect (> 2 SEM). Therefore, within the linear mixed-effects modelling procedures described below, CPM was included as a continuous predictor to capture the full range of individual variability in endogenous inhibitory control rather than imposing arbitrary categories. To avoid issues of collinearity and the false assumptions that can arise from re-modelling the same underlying variance, PDT was selected as the main predictor variable from among the three CPM measures as it contained the highest proportion of responders.

#### TSP analysis

It was initially intended that TSP would be calculated through the summation of normalised VAS measurements corresponding to each stimulus. However, recent publications have suggested that the statistical analysis of TSP paradigms is often hindered by baseline-dependant biases when simple first-minus-last comparisons are used, which can lead to ceiling effects or skewed directionality ^38^ and recommend assessing absolute change ^39^. Therefore, as magnitude of change is the primary variable of interest for TSP, and to attenuate baseline-dependant biases, the TSP effect delta score (Δ) was calculated as the mean of the VAS scores for the last three stimuli minus the first three stimuli representing an early-versus-late contrast. The applicability of this method is presented in Supplementary Figure 1.

Given the variability in TSP demonstrated through intra-class correlation coefficients (Supplementary Table 1), TSP responder rates were categorised based upon the direction and degree of change relative to standard error of measurement (SEM) as follows: Values < 1 SEM are weak responders; values between 1 SEM and 2 SEM are responders; values > 2 SEM are strong responders ^37^. As is typical, not all participants exhibited a strong facilitatory TSP effect (> 2 SEM). Therefore, within the linear mixed-effects modelling procedures described below, TSP was included as a continuous predictor to capture the full range of individual variability rather than imposing arbitrary categories.

### State-Trait Anxiety Inventory

The State-Trait Anxiety Inventory (STAI) was completed at the beginning of each session prior to cuff-algometry assessments. This psychometric questionnaire involves a twenty-item scale for two components, differentiating between the temporary condition of state anxiety and the general characteristic of trait anxiety ^40^.

Since its development, previous research has continuously validated this inventory in a range of populations, alongside confirming internal consistency and the separation of state-trait phenomenon ^41^. However, initial assessments of state and trait anxiety revealed a high correlation (*r* = .76, *t* (1, 273) = 19.44, *p* < .001) which remained functionally identical across sessions (Supplementary Figure 2). Alongside high collinearity (VIF > 49.5). Therefore, state and trait scores were averaged to represent overall within- and between-participant differences in anxiety.

### Data analysis

All statistical analyses were conducted using R (version 4.3.1; R Core Team, 2023a), a full list of packages and their specific use cases is included in Supplementary Table 2.

### Intraclass correlation coefficients

For all applicable measures, test-retest reliability was assessed through intraclass correlation coefficients (ICC 3,1), using a two-way mixed-effects model with absolute agreement for single rater measurements ^42^. ICCs were interpreted as follows: Values less than 0.5 were indicative of poor reliability; values between 0.5 and 0.75 were indicative of moderate reliability; values between 0.75 and 0.9 were indicative of good reliability; values greater than 0.9 were indicative of excellent reliability ^42^.

### Linear mixed-effects modelling

A linear mixed-effects modelling approach was applied to investigate changes in reported pain intensity over time, evaluate the consistency of these changes across sessions, and assess the influence of age, gender, anxiety, HFS pain intensity, CPM, and TSP. Visual inspections of normality, linearity and homoscedasticity were also performed.

Reported pain intensity (*Y*) was evaluated separately for each modality, with fixed effects for time (*β_1_*), session (*β_2_*), arm (*β_3_*), and their interaction (*β_4_*), alongside random intercepts for each individual (*u_i_*) to account for a within-subjects repeated-measures design. This model can be expressed as:

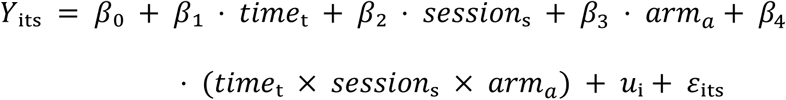

Visual inspections initially indicated that all reported pain intensity measures failed to meet assumptions, therefore, log-plus-one transformations were applied to stabilise variance and reduce deviations from linearity (Supplementary Figure 3a-b). Critically, this transformation preserved monotonicity and did not standardise variance.

Random slopes for time and session were also initially included to evaluate consistency issues within the repeated-measures design. However, within these models slope variance was estimated at or near zero, indicating singularity. Consequently, these did not improve model fit and were therefore removed, although comparisons are presented within Supplementary Figure 3c-d.

Within each successfully fitted model, conditional and marginal variance statistics were calculated using a mixed model R^2^, and partial ETA squared was used to estimate fixed effect size. The significance of fixed effects was evaluated via a three-way within-subjects Type III Analysis of Variance, and Satterthwaite’s method was applied to adjust degrees of freedom and to account for any uncertainty introduced by random effects and model complexity.

Where applicable, pairwise comparisons were performed by calculating estimated marginal means based upon the fixed and random effects of each model. Estimates were conducted separately per-session, with Dunnett’s adjustment for multiple comparisons applied to contrasts between baseline and each time point post-HFS. The resulting estimates were back-transformed through exponential-minus-one and standardised effect sizes are also reported.

Furthermore, to assess the effects of anxiety, CPM, TSP, age, gender, and pain intensity during HFS, each was added as a covariate (*Σβₖ*). Notably, although interactions between each covariate (*β_4_*) and the main effects (*β_1_*, *β_2_*, *β_3_*) were modelled, interactions between covariates were not run to avoid overfitting.

This can be expressed as:

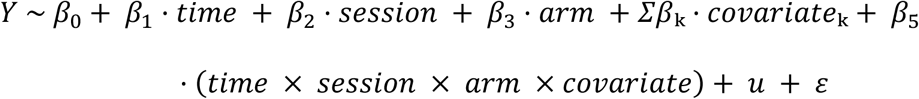

Additionally, due to errors during data collection two participants did not complete cuff-algometry testing. Therefore, the TSP and CPM covariates were assessed separately to avoid reducing the sample size when unnecessary (N = 27, (f = 17). All models were graphically assessed, resulting in the use of logarithmic-plus-one transformations as previously (Supplementary Figure 3e-h). Prior to modelling, numerical covariate measures were centred to support meaningful interpretations at intercept.

## Results

### Demographic characteristics

A total of 35 healthy adults were recruited to the study, however, the final sample included was 28, mainly due to feasibility issues during data collection as detailed within the OSF amendments (10 male: 32.2 ± 17.1 years [range 20 - 66]; 19 female: 27.6 ± 11.6 years [range 19 – 60]).

Exclusion criteria resulted in the removal of one participant: no history of neurological conditions, long-term health conditions, chronic pain, psychiatric conditions, steroid use, inflammatory disease, or immunosuppression; no allergies or severe skin conditions that may affect the methods or areas of testing; not currently pregnant.

### Mechanical pinprick sensitivity – Test-retest reliability

For the accurate characterisation of HFS-induced SHA it is critical that pinprick stimuli are perceived as painful at baseline ^1^. Within the present study cohort this was the case for all participants as demonstrated by Figure 3a.

**Figure 3.**
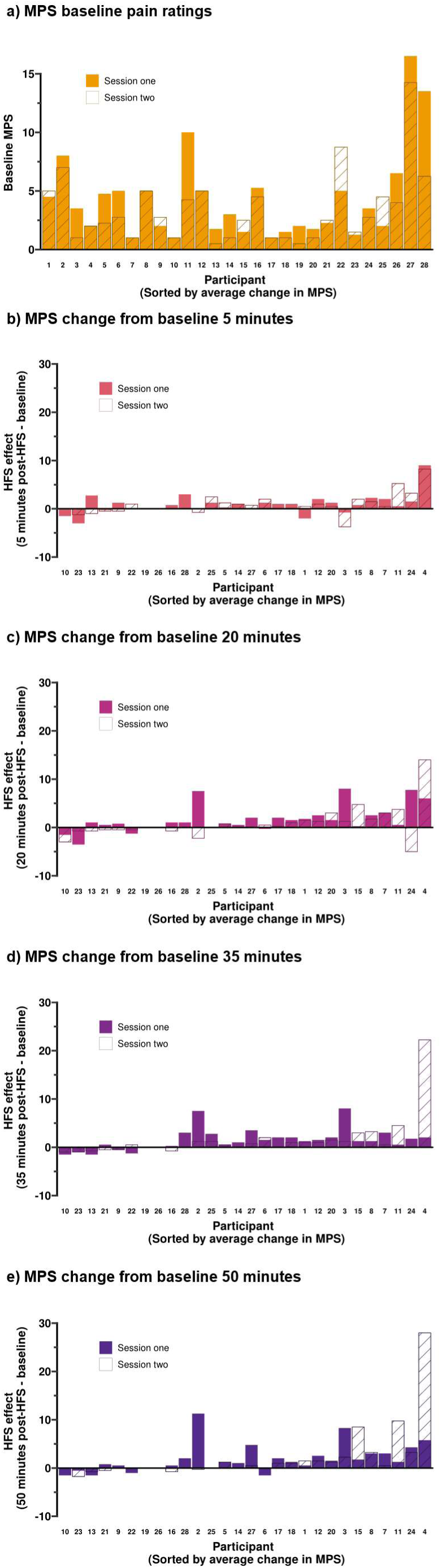
a) Test arm MPS baseline pain ratings and b-e) MPS change-from-baseline post-HFS. Individual variability plots were sorted by change-from-baseline in test arm MPS, which was averaged over all post-HFS time points and sessions for these rankings. This ordering is retained throughout Figure 3 and the relevant aspects of Figure 4 to aid interpretability. MPS = Mechanical pinprick sensitivity.

Furthermore, reliability assessments via intraclass correlation coefficients (ICC) are reported in Table 1 and indicate good reliability for MPS at baseline for both test (0.8) and control (0.79) arms. However, when assessing test arm MPS change from baseline, reliability across the four post-HFS time points was decreased (ICCs in ascending time order: 0.64, 0.15, 0.11, and 0.25) as demonstrated by Figure 3b-e.

### Mechanical pinprick sensitivity – Mixed effects modelling

The mixed-effects modelling reported in Table 2 explained 77% of the variance in pain ratings. However, fixed effects only accounted for 6%, suggesting that individual differences played a large role in the variability of reported pain intensity. Visual inspection demonstrated a positive influence of time when compared to pre-HFS for the test arm (Figure 2a). However, this was absent for the control arm. Statistical analyses revealed significant effects for time (*F* (4, 532) = 2.9, *p* = .02), session (*F* (1, 532) = 87.6, *p* = <.001), and arm (*F* (1, 532) = 34.8, *p* = <.001). Whilst only the interaction between time and arm was significant (*F* (4, 532) = 6.07, *p* = <.001). Therefore, when interpreted whilst considering the prior log-transformation, a multiplicative increase in pain intensity is evident following HFS induction within session 1. However, pain intensity was significantly reduced within session 2 in a manner that was consistent across both time and arms.

Within both sessions, pairwise comparisons to baseline of the estimated marginal means revealed that a statistically significant increase in pain intensity occurred for the test arm at 50 minutes post-HFS (session 1: 0.28 [0.08, 0.48], *p* = .004; session 2: 0.86 [0.31, 1.41], *p* = .01), whilst the control arm revealed no significant changes from baseline at any time point. Furthermore, although no aggregate time:session interactions were present, the time course of secondary hyperalgesia development differed between sessions: significantly increased pain sensitivity was present from 20 minutes post-HFS onwards during session 1 (20 mins: 0.25 [0.05, 0.45], *p* = .01; 35 mins: 0.27 [0.07, 0.46], *p* = 0.007), but only at 50 minutes post-HFS during session 2, as reported above.

### Single-pulse electrical pain perception – Test-retest reliability

Reliability assessments of single-pulse EPP pain ratings via intraclass correlation coefficients (ICC) are reported in Table 1 and indicate good-to-excellent reliability at baseline for the test (.92) and control (.80) arms.

### Single-pulse electrical pain perception – Mixed effects modelling

The mixed-effects modelling reported in Table 3 explained 86% of the variance in pain ratings. However, fixed effects only accounted for 4%, suggesting that individual differences played a large role in the variability of reported pain intensity. Visual inspection demonstrated no influence of time when compared post-HFS for the test arm (Fig. 2b). However, a slight deviation was present at 35 minutes for the control arm. Statistical analyses revealed no significant effects for time (*F* (4, 532) = 1.4, *p* = .23) or arm (*F* (1, 532) = .06, *p* = .81). Although, a significant effect of session was present (*F* (1, 532) = 137.5, *p* < 0.001). There were no significant interaction effects.

Pairwise comparisons to baseline of the estimated marginal means revealed no statistically significant changes in pain intensity for either the test or control arms. Therefore, no facilitation of the single-pulse electrical stimuli is evident following HFS induction.

### Influence of high-frequency stimulation pain intensity during conditioning

Reliability assessments of average pain during HFS conditioning were conducted via ICC, demonstrating good reliability (0.87 [0.73 0.94]) (Supplementary Table 1), which was supported by the visual inspection of individual variability (Figure 4a).

**Figure 4.**
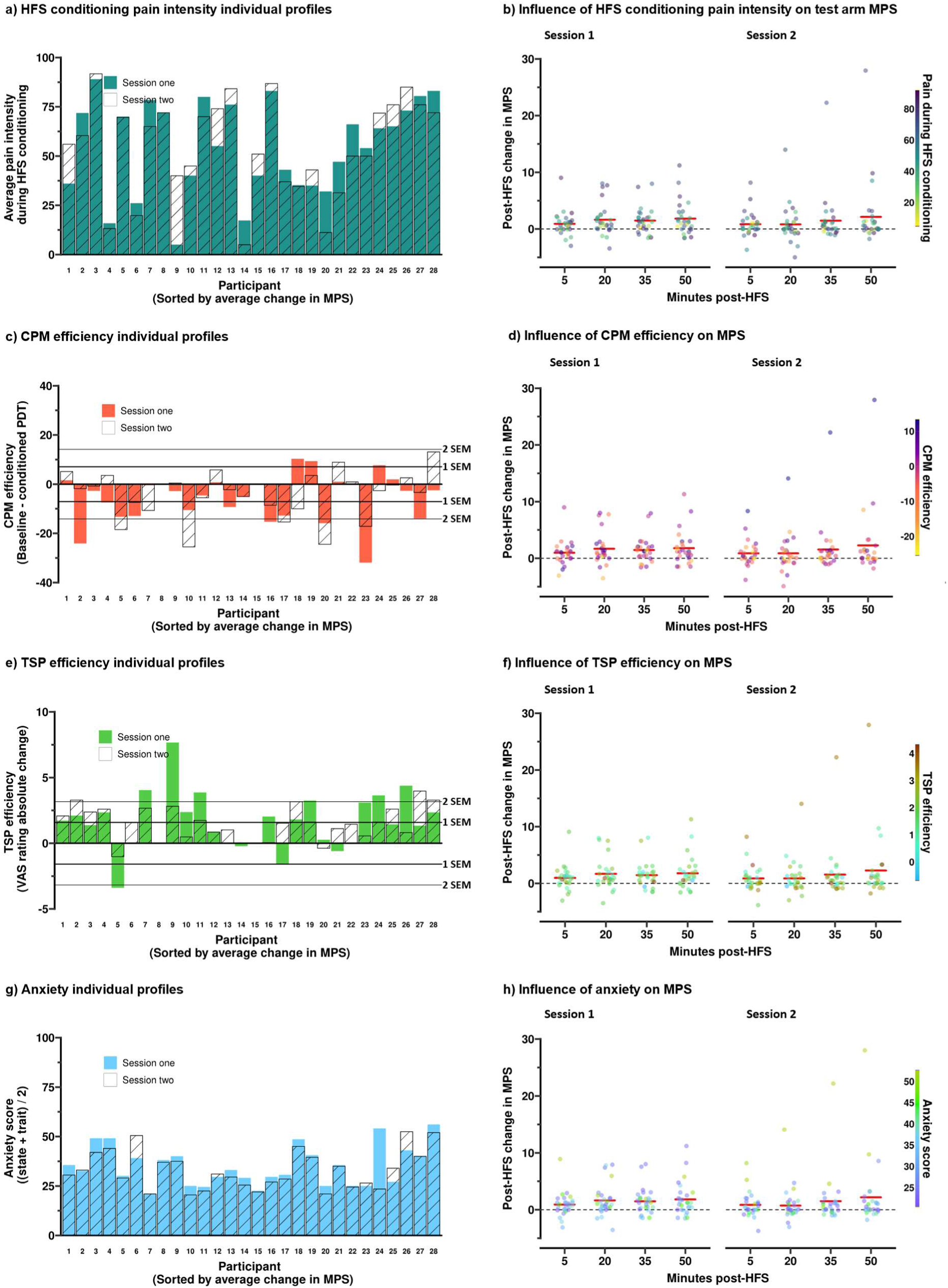
Individual variability plots (left) for pain intensity during HFS conditioning, anxiety, CPM, and TSP, alongside their respective influences on change in test arm MPS post-HFS. Factors: a-b) Average pain intensity during HFS conditioning; c-d) Pain detection threshold CPM efficiency; e-f) TSP efficiency; g-h) average anxiety scores. Individual variability plots are sorted by change-from-baseline in test arm MPS, which was averaged over all post-HFS time points and sessions for these rankings. This ordering is retained throughout Figure 4 and is identical to Figure 3 to aid interpretability. MPS scores are displayed as absolute change from baseline. Change in MPS over time was not significantly influenced by any factor (Tables. 3-4). MPS = Mechanical pinprick sensitivity; HFS = High-frequency stimulation; CPM = Conditioned pain modulation; TSP = Temporal summation of pain.

The inclusion of pain during HFS conditioning within the mixed-effects models resulted in an increase to fixed-effects explanatory power for both MPS (conditional *R*^2^ = .77, marginal R^2^ = .16) and EPP (conditional *R*^2^ = .84, marginal R^2^ = .13). When modelled across all time points, arms, and sessions, pain intensity during HFS conditioning demonstrated a significant positive influence on both MPS (F (1, 300.48) = 13.26, *p* < .001) and EPP (F (1, 417.59) = 17.99, *p* < .001) as reported in Table 4. However, whilst there were no significant interactions between pain during HFS and changes in MPS over time, across arms, or between sessions (all F < 1.78, all *p* > .17), there was a significant interaction between HFS pain intensity and how EPP changed over time (F (4, 519.38) = 3.11, *p* = .015). However, due to the lack of any significant changes in EPP over time, interpretation of this interaction is limited. Therefore, whilst higher pain during HFS predicted higher overall pain for both MPS and EPP, this did not explain inter-individual variability in the magnitude of post-HFS SHA development (Table. 4; Figure 4b), and although HFS pain intensity may have influenced homotopic plasticity, the lack of significant PHA does not allow for nuanced interpretation.

### Influence of Conditioned Pain Modulation

Reliability assessments of CPM were conducted via ICC, demonstrating moderate reliability (0.5 [0.15 0.74]) (Supplementary Table 1), which was supported by the visual inspection of individual variability (Fig. 4c).

The inclusion of CPM within the mixed-effects models resulted in an increase to fixed-effects explanatory power for both MPS (conditional *R*^2^ = .78, marginal R^2^ = .09) and EPP (conditional *R*^2^ = .86, marginal R^2^ = .05). When modelled across all time points, arms, and sessions, CPM demonstrated a significant positive influence on MPS (F (1, 519) = 7.33, *p* < .007), as reported in Table 5. However, there was no significant influence on EPP (F (1, 512) = 7.33, *p* < .24). Furthermore, there were no significant interactions between CPM and changes in MPS or EPP over time, across arms, or between sessions (all F < 2.22, all *p* = .14), as demonstrated by Figure 4d and reported in Table 5. Therefore, although CPM efficiency assessed at baseline predicted overall MPS pain intensity both at baseline and post-HFS, it did not predict inter-individual variability in SHA.

### Influence of Temporal Summation of Pain

Reliability assessments of TSP were conducted via ICC, demonstrating poor reliability (0.38 [0 0.66]) (Supplementary Table 1), which was supported by the visual inspection of individual variability (Figure 4e).

The inclusion of TSP within the mixed-effects models resulted in an increase to fixed-effects explanatory power for both MPS (conditional *R*^2^ = .79, marginal R^2^ = .08) and EPP (conditional *R*^2^ = .86, marginal R^2^ = .05). However, when modelled across all time points, arms, and sessions, TSP demonstrated no significant influence on MPS or EPP (all F < 1.17, all *p* > .28), as reported in Table 5. Furthermore, there were no significant interactions between TSP and changes in either MPS or EPP over time, session, or arm (all F < 2.17, all *p* > .17), as demonstrated by Figure 4f and reported in Table 5. Therefore, TSP assessed at baseline did not predict the development of HFS-induced secondary hyperalgesia in either session.

### Influence of anxiety

Reliability assessments of average anxiety were conducted via ICC, demonstrating good reliability (0.75 [0.52 0.87]) (Supplementary Table 1), which was supported by the visual inspection of individual variability (Figure 4g).

The inclusion of anxiety within the mixed-effects models resulted in an increase to fixed-effects explanatory power for both MPS (conditional *R*^2^ = .78, marginal R^2^ = .08) and EPP (conditional *R*^2^ = .87, marginal R^2^ = .05). When modelled across all time points, arms, and sessions, anxiety demonstrated no significant influence on MPS or EPP (all F < 0.2, all *p* > .66) as reported in Table 4. Furthermore, whilst there was no significant interaction between anxiety and changes in MPS over time or across sessions (all F < 1.01, all *p* > .4), as demonstrated by Figure 4h and reported in Table 4, there was a significant interaction between anxiety and arm (F (1, 521) = 6.32, *p* = .012). Therefore, anxiety may have exerted a lateralised influence over MPS but did not predict the magnitude of SHA development.

### Influence of age

The inclusion of age within the mixed-effects models resulted in an increase to fixed-effects explanatory power for both MPS (conditional *R*^2^ = .78, marginal R^2^ = .09) and EPP (conditional *R*^2^ = .86, marginal R^2^ = .09). When modelled across all time points, arms, and sessions, age demonstrated no significant influence on MPS or EPP (all F < 2.51, all *p* > .13) as reported in Table 4. Additionally, there were no significant interactions between age and changes over time or arm for either modality (all F < 3, all *p* > 0.8). However, there was a significant interaction between age and session for both MPS (F (1, 536.26) = 15.69, *p* < .001) and EPP (F (1, 531.57) = 19.37, *p* < .001). Subsequently, Johnson-Neyman visualisation revealed that changes in reported pain intensity between sessions were only significant for younger participants (<∼50 years old for both modalities) (Supplementary Figure 4). Therefore, whilst age did not explain inter-individual variability in the magnitude of SHA development post-HFS, it did predict overall variability in pain ratings across sessions.

### Influence of sex

The inclusion of sex within the mixed-effects models resulted in an increase to fixed-effects explanatory power for both MPS (conditional *R*^2^ = .78, marginal R^2^ = .08) and EPP (conditional *R*^2^ = .87, marginal R^2^ = .07). When modelled across all time points, arms, and sessions, sex demonstrated no significant influence on MPS or EPP (all F < 0.66, all *p* > .43) as reported in Table 4. Additionally, there were no significant interactions between sex and changes over time or arm for either modality (all F < 3.41, all *p* > .07). However, there was a significant interaction between sex and session for MPS (F (1, 536.26) = 15.69, *p* < .001), demonstrating that male participants experienced a significantly greater decrease in MPS across sessions (*η2* = −5.6 [−8.2, −3], *p* < .001) when compared to female participants (*η2* = −3.1 [−3.1, −4.5], *p* < .001). Therefore, whilst sex did not explain the inter-individual variability in the magnitude of SHA development post-HFS, it did predict overall variability in pain ratings across sessions.

## Discussion

The present study investigated the temporal dynamics of HFS-induced SHA between repeated test sessions. Test arm MPS measured in a heterotopic zone was significantly increased at 50 minutes post-HFS within both sessions, confirming the late-phase expression of SHA. However, whilst this facilitation was present from 20 minutes post-HFS during session 1, significant changes in MPS during session 2 were not evident during earlier time points (i.e., < 50 minutes). Therefore, a distinction between sessions in the early temporal dynamics of SHA development was present, which was reflected by poor test-retest reliability when change from baseline was assessed. In addition, these data show that CPM, TSP and anxiety had no relationship with the development of SHA and therefore further research into the factors influencing habituation across sessions is needed.

The current study has demonstrated that the trajectory of MPS change over time was not systematically altered between sessions, however there was a distinct reduction in the magnitude and time course of sensitivity development between sessions, indicating altered early temporal dynamics which may have been underpinned by an adaptive response in the processing and appraisal of repeated noxious stimuli following HFS conditioning across repeated test sessions ^43^.

Furthermore, in general, participants experienced lower evoked pain responses (i.e., MPS and EPP ratings both pre- and post-HFS in session 2 than in session 1. As this effect was present prior to HFS induction, it may reflect a general habituation to painful stimuli within session 2 ^44, 45^. Additionally, across both modalities, the degree of this reduction was dependent upon age and gender, demonstrating that older participants experience less variability in pain perception, whilst male participants experienced a greater reduction across sessions. However, whilst previous studies suggest that between-session habituation does not confound within-session sensitisation, increased pain thresholds often accompany such patterns ^46^. Therefore, as the present study applied identical HFS procedures within subjects, changes to the early temporal dynamics of secondary hyperalgesia development may be influenced by increased pain thresholds. However, as a statistically significant increase in sensitivity was present at fifty minutes in both sessions, irrespective of the overall reduced pain intensity and delayed onset in session 2, the consistent late-phase expression of mechanical pinprick hyperalgesia is supported across both sessions. Nonetheless, these findings reinforce the importance of carefully counterbalancing experimental conditions or the use of between-subject designs if HFS procedures are to be applied for interventional testing.

The absolute delivery intensity of both HFS and single-pulse electrical stimuli was chosen during session 1 based upon individual electrical detection threshold (EDT) and remained consistent throughout sessions for comparability. Previous studies have shown that EDT is lower in session 2 and that the intensity used to deliver HFS does not correlate with the magnitude of mechanical SHA across repeated test sessions ^9^. Indeed, a growing number of studies apply 3 mA HFS as a conditioning stimulus regardless of EDT ^17, 47^. In the present study, it is therefore unlikely that the reduced SHA response in session 2 is due to sub-optimal HFS conditioning, which we show produced comparable HFS pain intensity ratings across sessions. It is therefore possible that the reduced SHA response in session 2 is due to habituation to the psychophysical measures taken during the post-HFS period or resulting from differences in central pain processing during session 2.

Multiple factors could help to further explain the differences in the development of SHA between sessions. The DPMS is thought to be governed by top-down cortical influences which could be driving differences in the levels of pain sensitivity seen between sessions ^48^. However, when measured via CPM, no interaction was found with the session-induced reductions in pain intensity, although an overall predictive influence was present. Therefore, the present results demonstrate that stronger endogenous inhibition, as characterised through increased pain detection thresholds during CPM, predicted lower overall MPS both at baseline and post-HFS. However, the null interactions between CPM and changes in MPS over time suggests that post-HFS changes in MPS were not associated with CPM efficiency. Recent translational research has demonstrated that the activation of a diffuse noxious inhibitory control pathway during the induction of HFS has no capacity to suppress the spread of secondary hyperalgesia, as sufficiently high frequency excitatory signalling escapes inhibitory control ^20^. Furthermore, recent evidence has demonstrated that HFS does not activate the endogenous modulatory system ^19^. The results of the present study therefore reinforce these conclusions by demonstrating that CPM measurements of DPMS functioning immediately prior to HFS-induction have no predictive power for assessing the time course of secondary hyperalgesia development.

Furthermore, ascending nociceptive facilitation, as measured by the TSP, was hypothesised to demonstrate a direct predictive relationship with the development of secondary hyperalgesia, as wind-up mechanisms in the dorsal horn are implicated in both TSP and HFS paradigms. However, despite facilitation being present in both TSP and MPS measurements, the two were not significantly related. Again, Patel et al., demonstrated a similar lack of interaction, suggesting that the recruitment of different amplification mechanisms may present a feasible justification, such as differences in the engagement of peripheral afferent pathways, as HFS targets transcutaneous nociceptors whilst cuff-algometry activates deep tissue afferents ^20^. The statistical analysis of TSP paradigms is often hindered by baseline-dependant biases, which can lead to ceiling effects or skewed directionality ^38^. Although the present study attempted to rectify this underlying methodological limitation, the poor TSP reliability and session-specific directionality demonstrated by the present results suggest that measurements of ascending facilitation evoked by cuff pressure may not be precise enough to accurately predict variations in MPS. Additionally, temporal summation of pain (TSP) was assessed through cuff algometry, a standard method reflecting the facilitatory capacity of ascending nociceptive pathways within the spinal cord ^48, 49^. However, whilst TSP provides a general measure of spinal excitability, cuff algometry primarily activates deep tissue nociceptors. Therefore, the lack of significant interactions between TSP and HFS-induced SHA may be due the engagement of different populations of nociceptive fibres.

The State-Trait Anxiety Index was also utilised to examine how anxiety may influence the development of HFS-induced SHA. The present findings suggest that the influence of anxiety was limited to lateralised differences in pain perception as anxiety was positively predictive of differences in MPS across arms, whilst there was no influence on the development of SHA. This conclusion is partly supported by previous evidence which has demonstrated how the consistent and prolonged experience of anxiety results in decreased pain thresholds and the amplification of acute pain perception ^18, 50^. The present findings suggest that the development of SHA is not mechanistically related to anxiety. However, the present study only assessed a healthy population, therefore, the relationship between long-term anxiety and SHA may be altered for individuals with clinically established symptoms. This contrasts with findings that show that pain expectations and the fear of pain can influence the magnitude of secondary hyperalgesia ^17, 47^. Additionally, the present study measured anxiety at baseline rather than induced anxious expectations of pain. Therefore, whilst experimental manipulation within clinically established anxiety disorders can produce neurological patterns that convincingly overlap with those found in phobic disorders, the degree of anxiety within the present study may have been too low for a significant influence ^51^.

Finally, the present results demonstrate a lack of substantial HFS-induced homotopic sensitivity changes in contrast with previous publications ^2, 4, 10, 11^. It is possible that within the area of homotopic plasticity, post-HFS changes are not sensitive to electrical stimulation. Therefore, future research is necessary to clarify the afferent fibres involved in HFS-induced homotopic plasticity. Klein et al., ^2^ and Pfau et al., ^4^ have previously demonstrated increased homotopic EPP post-HFS when compared to control arms, first onset at ∼20 minutes and peaking at ∼40 minutes. Alternatively, recent replications by van den Broeke et al., ^10, 11^ have consistently reported a reduction in homotopic EPP for both the test and control arms post-HFS. However, although they partially replicated comparatively increased test-verses-control arm EPP within one study ^11^ the opposite effect was observed within the second study ^10^. van den Broeke et al., ^10^ suggest that this disparity arises from differences in HFS-induction intensity. Although not completely analogous, the present results provide evidence supporting the conclusion that the intensity of HFS influences homotopic EPP, as assessed through the pain reported during HFS conditioning, which significantly predicted changes in homotopic EPP.

In conclusion, these data have provided test-retest temporal dynamics of HFS-induced changes in pinprick sensitivity across repeated test sessions with a distinct reduction in the magnitude and early temporal dynamics of sensitivity development during session 2. Whilst pain during HFS conditioning, anxiety, CPM, and TSP demonstrated various relationships with general mechanical pinprick pain intensity across all time points, arms and sessions, none of these measures specifically influenced the time course of secondary hyperalgesia development, suggesting that these measures were inadequate to explain the between-sessions variance in the SHA development time course. Therefore, these results suggest that careful consideration into experimental design should be applied in future studies investigating the development of HFS-induced SHA and further research into the factors influencing habituation across sessions is needed.

## Disclosures

This work is funded through a PhD studentship from the University of Plymouth (School of Psychology) and an Academy of Medical Sciences Springboard Award (Hughes: SBF007\100108). The authors have no conflicts of interest.

## Authorship Contributions

SM: Conceptualisation, funding acquisition, data collection, formal analysis, writing, reviewing and editing.

GG: Conceptualisation, funding acquisition, supervision, methodology, reviewing and editing.

SH: Conceptualisation, funding acquisition, supervision, methodology, writing, reviewing and editing, project administration.

## Data availability statement

Data are available upon request.

## Notes

### Competing Interest Statement

The authors have declared no competing interest.

