## Supplementary Information for "Temporal changes in mechanical pin prick sensitivity following high frequency induced sensitisation of central nociceptive pathways: a test re-test reliability study"

Supplementary Table 1

**Summary of covariate reliability assessments.**

| **Intraclass correlation coefficients**: 3, 1 [95% confidence intervals] | |
|---|---|
| Measure | Coefficients |
| Conditioned pain modulation (PDT) | 0.5 [0.15 0.74] |
| Temporal summation of pain | 0.38 [0 0.66] |
| Average anxiety | 0.75 [0.52 0.87] |
| Pain during HFS conditioning | 0.87 [0.73 0.94] |

**Supplementary Table 2. Summary of covariate reliability assessments**. Intraclass correlation coefficients were calculated using a two-way mixed-effects model with absolute agreement for single rater measurements (3,1). Coefficients and confidence intervals were computed independently for each time point, arm, and stimulation modality. Values less than 0.5 are indicative of poor reliability, values between 0.5 and 0.75 indicate moderate reliability, values between 0.75 and 0.9 indicate good reliability, and values greater than 0.9 indicate excellent reliability. HFS = high frequency stimulation; PDT = Pain detection threshold.

Supplementary table 2

**R Packages used for statistical analysis**

| Package | Use Case | Citation |
|---|---|---|
| Tidyverse | Data importing and manipulation. ggplot2 plotting features. | Wickham H, Averick M, Bryan J, Chang W, McGowan LD, François R, Grolemund G, Hayes A, Henry L, Hester J, Kuhn M, Pedersen TL, Miller E, Bache SM, Müller K, Ooms J, Robinson D, Seidel DP, Spinu V, Takahashi K, Vaughan D, Wilke C, Woo K, Yutani H. Welcome to the tidyverse. *J Open Source Softw* 2019;4(43):1686. |
| Magrittr | Pipe operators for in-place transformations. | Bache SM, Wickham H. magrittr: A Forward-Pipe Operator for R. R package version 2.0.3. 2014. Available from: https://CRAN.R-project.org/package=magrittr |
| Flexplot | Visualisation of data distributions. | Fife DA. flexplot: Graphically Based Data Analysis Using flexplot. *J Open Source Softw* 2020;5(52):2381. |
| Flextable | Reporting of model outputs. | Gohel D, Skintzos P. flextable: Functions for Tabular Reporting. R package version 0.9.2. 2023. Available from: https://CRAN.R-project.org/package=flextable |
| Lme4 | Fitting linear mixed-effects models (lmer()). | Bates D, Mächler M, Bolker B, Walker S. Fitting linear mixed-effects models using lme4. *J Stat Softw* 2015;67(1):1–48. |
| LmerTest | Improved lmer() model outputs. | Kuznetsova A, Brockhoff PB, Christensen RHB. lmerTest package: tests in linear mixed effects models. *J Stat Softw* 2017;82(13):1–26. |
| Performance | Visualisation of model fits. | Lüdecke D, Ben-Shachar MS, Patil I, Waggoner P, Makowski D. performance: Assessment of regression models performance. CRAN. 2021. Available from: https://CRAN.R-project.org/package=performance |

| | | |
|---|---|---|
| Effectsize | Calculation of modelled effect sizes. | Ben-Shachar MS, Lüdecke D, Makowski D. effectsize: Estimation of effect size indices and standardized parameters. *J Open Source Softw* 2020;5(56):2815. |
| Emmeans | Calculation of estimated marginal means, contrasts, and effect sizes. | Lenth RV. emmeans: Estimated marginal means, aka least-squares means. R package version 1.8.8. 2023. Available from: https://CRAN.R-project.org/package=emmeans |
| Patchwork | Improved multi-plot visualisation. | Pedersen TL. patchwork: The Composer of ggplots. R package version 1.1.2. 2022. Available from: https://CRAN.R-project.org/package=patchwork |
| ggprism | Improved plot themes. | Mackinnon SP. ggprism: A ggplot2 extension inspired by GraphPad Prism. R package version 1.0.2. 2021. Available from: https://CRAN.R-project.org/package=ggprism |
| Interactions | Johnson-Neyman plots for visualisation of 2-way interactions. | Long JA. interactions: Comprehensive, User-Friendly Toolkit for Probing Interactions. R package version 1.2.0. 2024. Available from: https://CRAN.R-project.org/package=interactions |

**Supplementary Table 3. R Packages used for statistical analysis.** A full list of packages and their specific use cases. All statistical analyses were conducted using R. R Core Team. R: A language and environment for statistical computing. Vienna, Austria: R Foundation for Statistical Computing; 2023. Available from: https://www.R-project.org/.

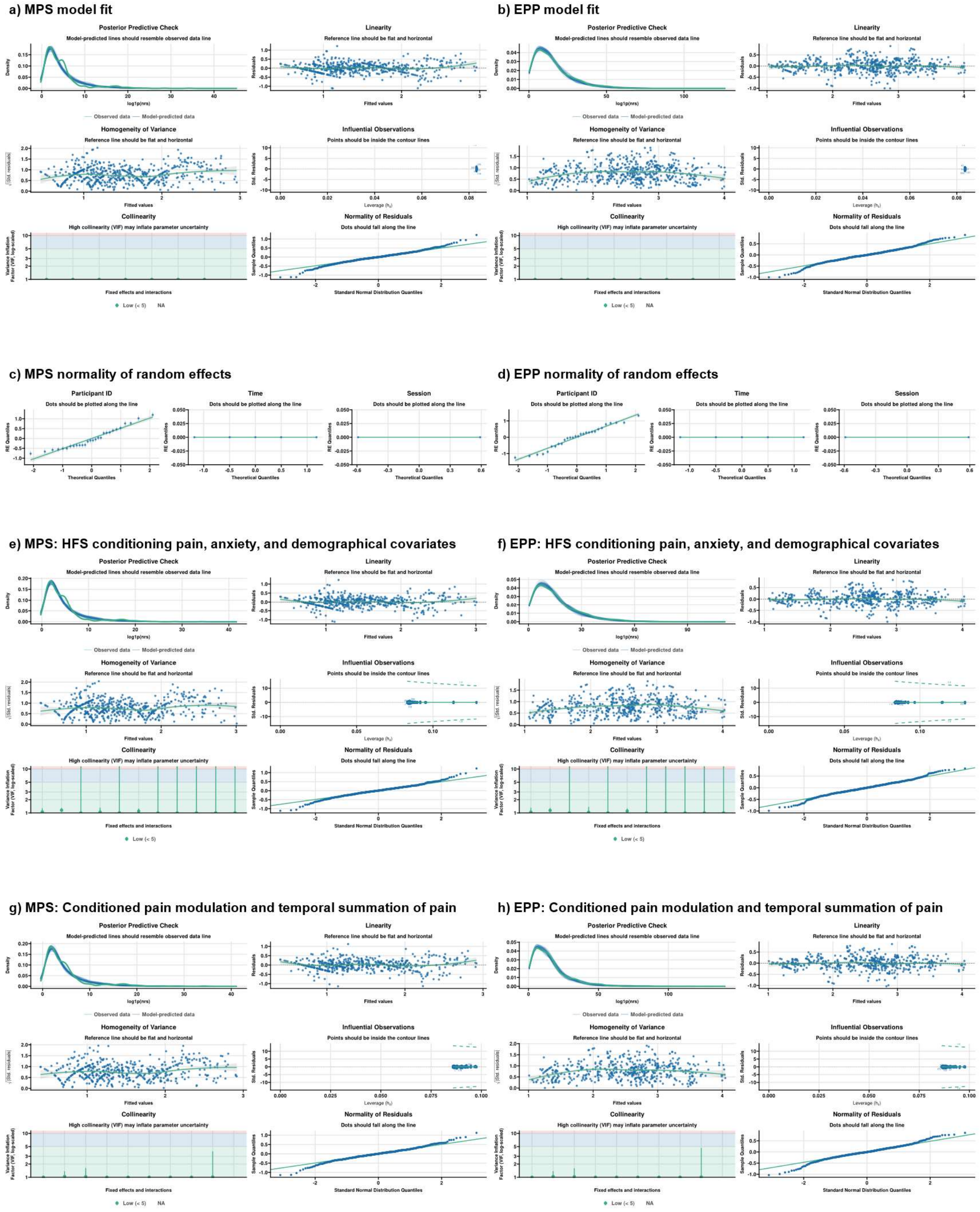


**Supplementary Figure 1. Visualisation of log1p transformed mixed-effects models for MPS (left) and EPP (right) displaying a-b) Model fits, alongside c-d) random effects slopes and e-h) models including covariates.** Model performance metrics were assessed using the Performance package in R (version 0.13.0; Lüdecke et al., 2021). MPS = Mechanical pinprick sensitivity; EPP = Electrical pain perception.

### TSP efficiency subject-level calculations

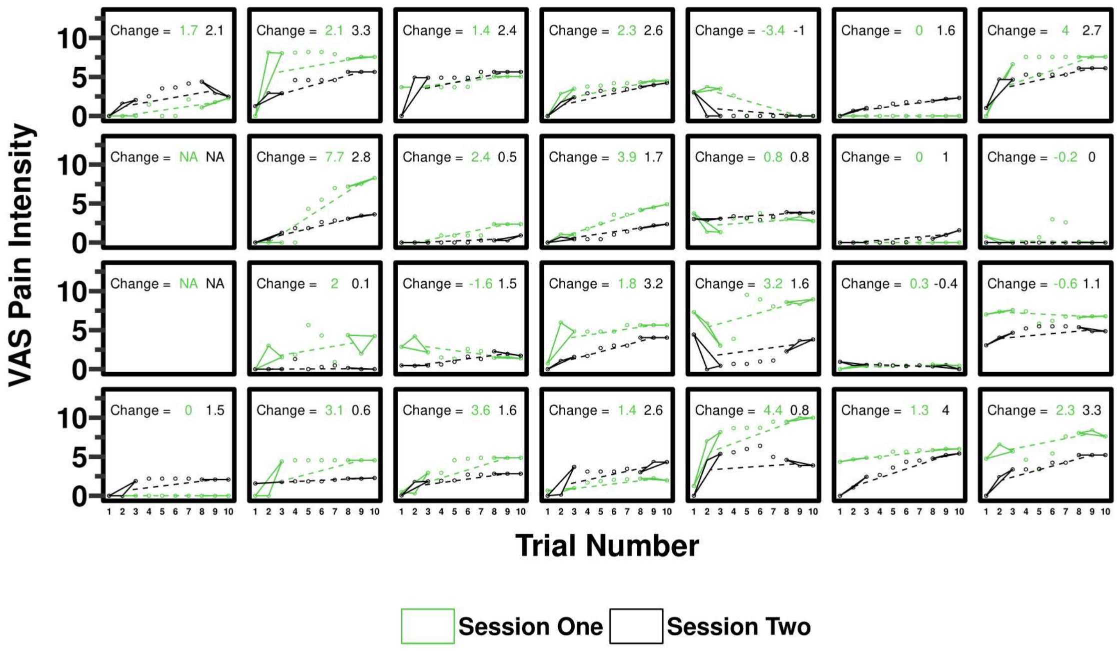


**Supplementary Figure 2. Subject-level absolute change (delta) calculations for the temporal summation of pain**. Absolute change between the first three and last three stimuli are displayed across sessions for each subject, sorted by average change in MPS. Due to an error during data collection, two participants did not complete TSP testing, therefore: N = 27, (f = 17).

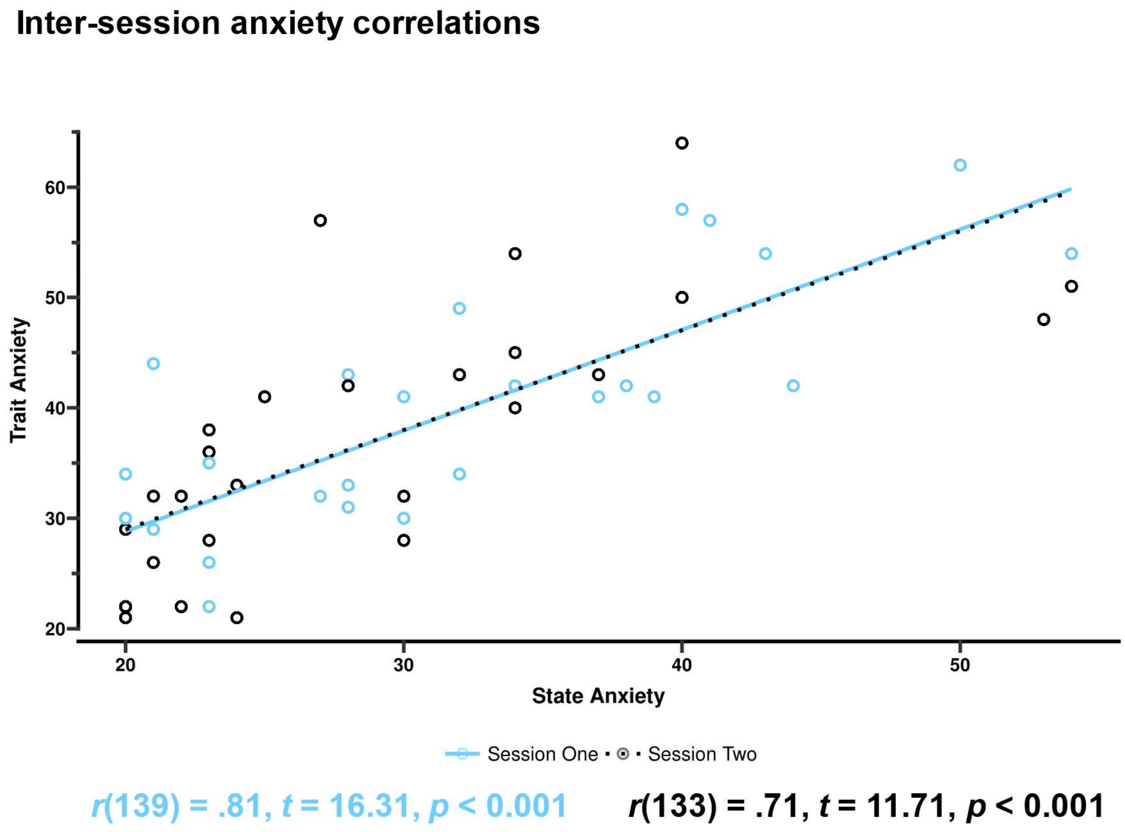


**Supplementary Figure 3. Correlations between state and trait anxiety within both sessions.** Due to an error during data collection, one participant did not complete the state questionnaire during session two.

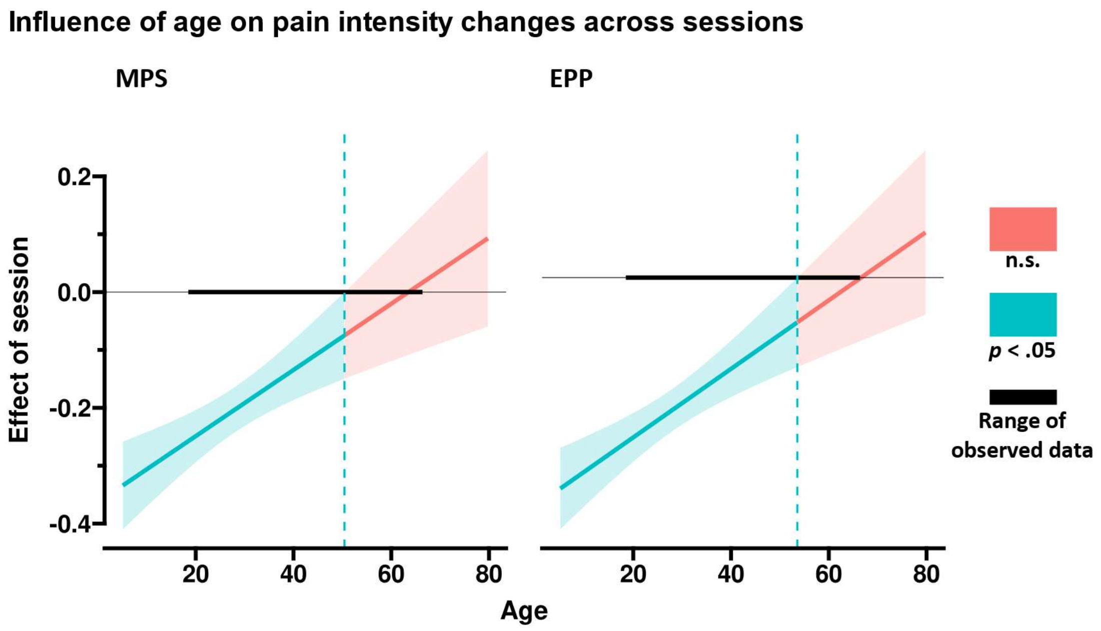


**Supplementary Figure 4. The influence of age on pain intensity changes across sessions for both MPS and EPP.** MPS = Mechanical pinprick sensitivity; EPP = electrical pain perception.
